# Phloem structure and development in *Illicium parviflorum*, a basal angiosperm shrub

**DOI:** 10.1101/326322

**Authors:** Juan M. Losada, N. Michele Holbrook

## Abstract

Recent studies in canopy-dominant trees revealed a structure-function scaling of the phloem. However, whether axial scaling is conserved in woody plants of the understory, the environments of most basal-grade angiosperms, remains mysterious. We used seedlings and adult plants of the shrub *Illicium parviflorum* to explore the anatomy and physiology of the phloem in their aerial parts, and possible changes through ontogeny. Adult plants maintain a similar proportion of phloem tissue across stem diameters, but scaling of conduit dimensions and number decreases the hydraulic resistance towards the base of the plant. Yet, the small sieve plate pores resulted in an overall higher sieve tube hydraulic resistance than has been reported in other woody angiosperms. Sieve elements scaled from minor to major leaf veins, but were shorter and narrower in petioles. The low carbon assimilation rates of seedlings and mature plants contrasted with a three-fold higher phloem sap velocity in seedlings, suggesting that phloem transport velocity is modulated through ontogeny. While the overall architecture of the phloem tissue in basal-angiosperm understory shrubs scales in a manner consistent with trees, modification of conduit connections may have allowed woody angiosperms to extend beyond their understory origins.

## INTRODUCTION

The ecological context in which angiosperms evolved has been hotly debated in the last decades. This is, in part, due to the apparent contradiction between the relative recent rise of angiosperms, measured in geological time (approximately 125 million years ago), and the rapid diversification that led to their ecological dominance (Feild *et al*., 2003a; 2004). The success of flowering plants correlates with the evolution of anatomical innovations that improved their physiological performance in different environmental scenarios, such as leaf reticulate venation and large xylem vessels (Zwieniecki & Boyce, 2014; Boyce & Lee, 2017). Observations of fossils from ancestral angiosperm lineages revealed that their anatomical features mirror those of living members of the basal grade, which include a small vessel lumen fraction in the stems (Bailey & Nast, 1948; Carlquist, 1982; Carlquist & Schneider, 2002), leaves with thick cuticles and large stomata, as well as irregular leaf venation (Hickey & Doyle, 1977; Upchurch *et al*., 1984; Carpenter, 2005; Coiffard *et al*. 2006). These data, combined with recent morpho-physiological measurements using extant taxa from the basal grade, point to wet, understory and disturbed habitats as a credible niche for the rise of angiosperms (Feild *et al*. 2003b, 2004; Feild & Arens, 2005, 2007; Barral *et al*., 2013). Most extant lineages of woody basal flowering plants are restricted to the understory areas of a few tropical environments. Understanding the structure and physiology of basal grade angiosperm phloem may shed light on growth constraints during their initial expansion in the Cretaceous, as well as bring information onto the hydraulic properties of the phloem of woody plants in the understory, so far overlooked.

A critical constraint of understory growth is the low availability of light and the consequent low rates of photosynthesis. Indeed, *in situ* measurements of the maximum photosynthetic activity in these conditions confirmed that low primary productivity is associated with a similarly low xylem hydraulic performance (Feild *et al*., 2005). While the hydraulics of the xylem are coupled with the sugar loading of the phloem in the mesophyll (Hölttä *et al*. 2006; Liesche *et al*. 2011; Nikinmaa *et al*. 2013; Rockwell *et al*., 2018), the performance of the phloem in leaves is still poorly understood. With a recent exception (Carvalho *et al*., 2017a), detailed evaluations of phloem architecture in leaves with reticulate venation are lacking. This is in part due to the challenging manipulations required to quantify the geometry of the sieve tubes in the tapering veins, as well as difficulties in evaluating the velocity of the sap inside the sieve tubes, typically measured by a fluorescent dye tracer (Jensen *et al*., 2011, Etxeberria *et al*., 2016) or radiolabeled compounds (Knoblauch *et al*., 2016). Thus far, phloem sap velocities in leaves (including petioles) have been measured in only a handful of adult plant species (Jensen *et al*., 2011), and in the seedlings of the family Cucurbitaceae (Savage *et al*., 2013). Surprisingly, despite the critical interplay between development and physiological performance of woody plants during their ontogeny, comparisons between seedlings and adult plants have been missing so far.

The widely accepted mechanism of phloem transport is the differential osmotic gradient generated between sources and sinks (Münch, 1930; Thompson & Holbrook, 2003; Pickard, 2012; Knoblauch & Peters, 2017), but the validity of this hypothesis has only recently been tested in herbaceous vines (Knoblauch *et al*., 2016), and trees (Liesche *et al*., 2017; Savage *et al*., 2017). In canopy dominant tree species, long distance geometrical scaling of the sieve elements of the phloem is conserved, resulting in a reduced resistivity of the individual collecting tubes towards the base of the tree (Savage *et al*., 2017). Yet, understanding the overall flow capacity of the phloem requires quantification of sieve tube numbers alongside vascular tapering (i.e. tree/shrub branching). Sieve tube quantification has only been explored at single points of the stems in a few trees (Lawton & Canny, 1970; Ghouse *et al*., 1976; Ghouse & Jamal, 1979; Khan *et al*., 1992), but never comparing different stem diameters, essential to evaluate whole plant transport.

Pursuing these questions, we investigated the angiosperm shrub *Illicium parviflorum* to gain a better understanding of phloem structure and function in the aerial parts of plants adapted to understory environments and through their ontogeny. Despite a restricted distributional range in the tropics of America and Asia, the family Illicaceae is the largest within Austrobaileyales, one of the three lineages that compose the basal grade of flowering plants, only predated by Nymphaeales and the monotypic Amborellales (Mathews & Donoghue, 1999; Parkinson *et al*., 1999; Qiu *et al*., 2000; Soltis *et al*., 1999; 2018). Little is known of the phloem in these lineages (Bailey & Nast, 1948) and this information could bring insights into their growth patterns and ecological adaptations. Our results include a novel quantification of sieve tube element number per cross sectional area of the stems and major veins of leaves, and characterization of phloem architectural traits related to the hydraulic efficiency of carbohydrate transport.

## MATERIALS AND METHODS

### Plant materials and growth conditions

Mature plants of *Illicium parviflorum* were grown in plastic pots in the greenhouses of the Arnold Arboretum of Harvard University, using high porosity growing medium PRO-MIX (PremierTech Horticulture and Agriculture Group, Riviere-du-Loup,QC, Canada), and under conditions of 25 °C ± 2°C and RH of 75% (Fig. 1). Plants were watered to field capacity every morning. Three seeds per pot were sown, and first seedlings (with two cotyledons) were visible four months after sowing. They were repotted individually and grown for three more months until the emergence of the first true leaves (approximately 3cm long), which were used for the measurements described below. Due to the understory habit of these plants and based on our previous observations, high sun exposure was avoided with a dark net between the greenhouse roof and the plants (average PAR 100 µmol m^−2^ s^−1^).

**Fig. 1.**
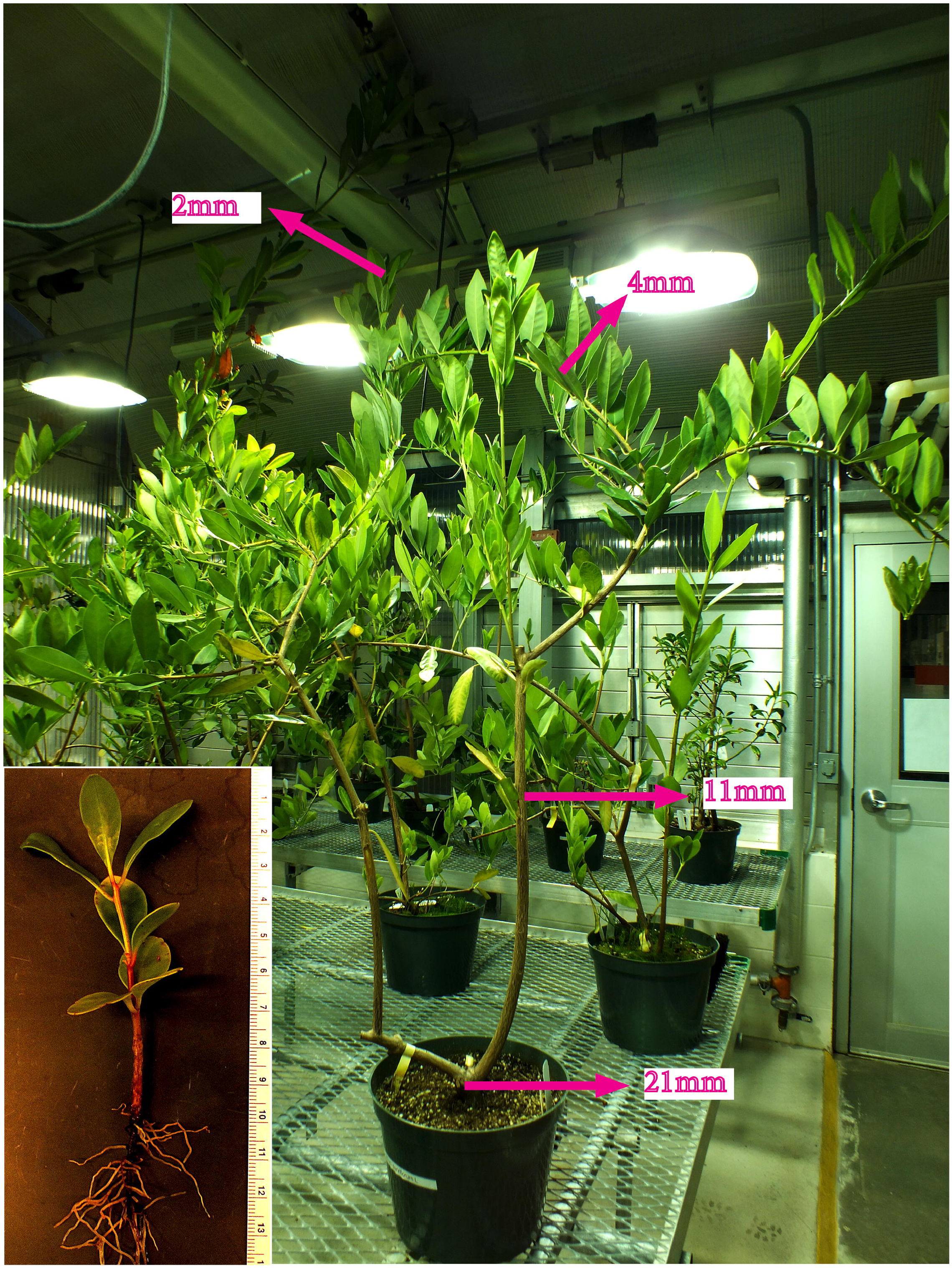
*Illicium parviflorum* shrubs in the greenhouse. Arrows indicate the diameter of the stem where samples were obtained. Distal parts correspond with thinner stems and areas of primary growth. Inset shows an image of an uprooted seedling. Scale in cm.

### Measurements of carbon assimilation

Gas exchange was measured in three exposed leaves from three branches oriented in all directions in each mature plant (n=3 adult plants), as well as two leaves from each seedling (n=4 seedlings). Measurements were done at three time points during the day (8:30am, 12:30am, and 4:30pm), once per week during February and March 2017. We used a LI-COR 6400 (with light source) configured to track ambient PAR conditions at each measurement (average 100 µmol m^−2^ s^−1^). The reference CO_2_ was 400 µmol mol^−1^, the average leaf temperature 20.0°C ± 0.16°C (SD), and vapor pressure deficit 1.06 ± 0.06 kPa (SD).

### Sample preparation for stem anatomy

Due to the shrubby nature of mature *Illicium parviflorum* plants, lateral branches were divided into four stem diameter classes (see Fig. 1): <2mm (primary growth), 4-6mm (green shoots with secondary growth), 9-11mm (stems with bark), and >21mm (main stem). The length of each branch segment was measured, and the number and area of the leaves per unit length evaluated (n=30). To observe changes in the vascular tissues with increasing stem diameters, 5-10cm long stem sections from each diameter class and three different plants were cut with clippers and kept in 1X Tris-buffered saline (TBS). 50μm thick cross-sections were then obtained with a Reichert-Jung Hn-40 sliding microtome (Austria), immediately mounted onto glass slides, and stained with a solution of 0.1% aniline blue PO_4_ K_3_ (Linskens & Esser, 1957), which stains callose of the sieve plates. Samples were observed with either a Zeiss Axiophot microscope with epifluorescence and an AxioCam 512 Color connected to the AxioVision software (Zeiss, Oberkoche, Germany), using the DAPI narrow filter band (excitation 365nm, bandpass 12 nm; dichroic mirror FT 395nm; barrier filter LP397nm). Individual images of the stem cross sections (taken with the 5x/0.15 Plan-Neofluar objective) were aligned and merged into a composite image to visualize and measure the areas of the whole stem with the Adobe Photoshop software (Adobe Systems Inc., Newton, MA, USA).

### Evaluation of sieve tube geometry in stems

To evaluate the length and the radius of the sieve tube elements, a second set of samples from the same stem diameters described above were hand-sectioned with a micro-scalpel, mounted on slides and stained with a mixture of 0.1% aniline blue in PO_4_ K_3_, and 0.1% calcofluor white in 10mM CHES buffer with 100mM KCl (pH=10), which stains the cellulose from the cell walls of the sieve tube elements, prior to microscopic observations.

In order to quantify the number of sieve tube elements, a third set of samples (2, 4, 6 and 11mm diameter) of about 0.5cm thickness were collected and then fixed in 4% acrolein (Polysciences Inc., Warrington, PA, USA) in a modified piperazine-N,N’-bis (2-ethanesulfonic acid) (PIPES) buffer adjusted to pH 6.8 (50 mM PIPES and 1 mM MgSO4from BDH, London, UK; and 5 mM EGTA from Research Organics Inc., Cleveland, OH,USA) for 24 hours, then rinsed thrice in the same buffer, and finally dehydrated through a series of increasing ethanol concentrations (10%, 30%, 50%, 70%, 80%, and 100%), one hour each. Samples were incubated in the catalyzed solution of the resin Technovit 8100 (Electron Microscopy Sciences, Hatfield, PA, USA) for at least three months, and finally embedded under anoxic conditions and at 4°C. Later, blocks were mounted in microtome studs and serially sectioned at 4μm with a Leica RM2155 rotary Microtome (Leica Microsystems Inc., Germany). After mounting them in Superfrost slides, they were stained with aniline blue and calcofluor as described previously to quantify both the number of sieve tube elements and the total number of cells per phloem sectional cluster (the areas of the phloem separated by ray parenchyma).

All samples were imaged with a Zeiss LSM700 Confocal Microscope (20x/0.8 M27 Plan-Apochromat objective for cross-sections and 63x/1.40 Oil DIC M27 Plan-Apochromat objective for longitudinal sections) using the 405nm laser band to excite the sample, and a Zen Black 2010 software connected to a Zeiss HR camera to create the final compound tiles (Zeiss, Oberkoche, Germany). Serial tile images obtained from resin sections were aligned with the Adobe Photoshop software (Adobe Systems Inc., Newton, MA, USA), and then both parenchyma cells as well as sieve tube elements quantified. To estimate the total number of sieve tubes per stem diameter, we calculated the tube number per cluster area, multiplied by the calculated area of the phloem at each stem diameter. Note that this will slightly overestimate the number of sieve tubes, as the phloem clusters are separated by typically uniseriate rays.

### Evaluation of sieve plate geometry

To evaluate in detail the size of pores that make up the sieve plates, another batch of wood sections were cut and immediately frozen in liquid nitrogen, transferred to super-chilled ethanol, and then sectioned in the same orientation of the sieve plates at each diameter. After that, the cut sections were incubated within a mixture of 0.1% proteinase K dissolved in 50mM Tris-HCl buffer, 1.5mM Ca^2+^ acetate and 8% Triton X-100, pH 8.0 (Mullendore *et al*., 2010), using a water bath at 60°C for a period of two weeks. After rinsing with ethanol to deactivate the proteinase and washing them thrice with water, they were incubated with a 1% aqueous solution of α-amylase for at least two days at 60°C, then rinsed thrice in water and finally freeze dried for 24 hours. Following, samples were mounted on SEM studs, and sputter coated with gold-palladium using a Denton Vacuum Desk II Sputter Coater for 180 secs at 20Volts and 50militor pressure. Studs with samples coated with gold-palladium were imaged with a JEOL-6010LV scanning electron microscopy (SEM) (JEOL Inc., Peabody, Massachusetts, USA) using high vacuum and an accelerating voltage of 10-15kV. Pore size was evaluated in at least ten samples (n=50 pores) of the largest stem diameters (11mm and 21mm).

### Sample preparation for leaf anatomy

Transverse hand sections of the petiole, the midrib, and the secondary veins of mature *I. parviflorum* leaves were serially obtained, stained with aniline blue and calcofluor, and finally imaged with a confocal microscope (details below). To understand changes in the area of the xylem, phloem and sclerenchyma along the major vein of mature leaves, three sequential cross-sections or the petiole, the midrib and the tip of the major vein were obtained from five different mature leaves, then imaged and manually outlined with the Image J software (Supporting Information Fig. S1). Longitudinal sections of the same vein areas from the petiole, the midrib, second and third order veins were obtained from five different leaves and imaged with the protocol described below for quantification of length and radius of the sieve tube elements.

Due to the small size of leaves from seedlings, leaf material fixed in 4% acrolein, followed by embedding in Technovit 8100 prior to serial sectioning with a microtome as previously described. Variation among areas, sieve tube lengths, radius, and numbers from leaves were averaged and means compared among the different vein orders using a one way ANOVA and Tukey test at a P<0.05. All statistical analysis were performed with SPSS software (SPSS Inc., Chicago, USA).

### Measurements of phloem sap velocity

Phloem transport velocity was measured by tracking the movement of the fluorescent dye carboxifluorescein (CF) (reviewed in Knoblauch *et al*., 2015), in the secondary veins of both living mature plants and seedlings at 11:00am each day. After saturating the soil with water, three mature plants and nine seedlings were taken to the lab and the leaves were immobilized to the microscope stage with the abaxial surface exposed upwards. A 10μL droplet of a mixture containing 0.01 M CF diacetate in 1:10 mixture of acetone and distilled, deionized water, supplemented with and 0.1% of the surfactant SilEnergy (RedRiver Specialties Inc., Shreveport, LA, USA) was applied with a micropipette to the distal most part of the secondary veins (adapted from Jensen *et al*., 2011, Savage *et al*., 2013). To allow a better permeabilization of the dye, the tip of the pipette was used to slightly abrade the thick cuticle, and the small window opened (approximately 1mm^2^) was covered with the liquid during the course of the experiment to prevent desiccation. To track the movement of the dye, we used a portable Stereo Microscope Fluorescence Adapter with 510-540nm excitation wavelength, and a long pass 600nm filter band (Nightsea, Lexington, MA, USA). Time-lapse images were obtained every ten seconds for a period that spanned from 20 to 40 minutes with a Zeiss v12 Dissecting microscope using the 0.63X PlanApo objective and an AxioCam 512 Color camera connected to the AxioVision software (Zeiss, Oberkoche, Germany). Velocity was calculated by tracking the time taken by the dye front to travel a known distance.

## RESULTS

### Anatomy of the stems in *Illicium parviflorum*

*Illicium parviflorum* shrubs have numerous lateral branches and a short thicker stem (Fig. 1). Younger branches are green (2-10mm diameter) and support spirally arranged leaves, whereas older branches (>10mm diameter) are barky and seldom retain leaves. The length of the branches scales proportionally with their diameter, and thus the stems with 2mm diameter reach up to 10cm long, whereas 6-10mm diameter stems extend through 50cm. While the ratio of leaf number to stem length is higher in the youngest stems (0.7±0.07SE) compared with older ones (0.5±0.03SE), the average leaf area increases from 13.4cm^2^ in the thinner stems (2mm diameter, primary growth) to 20.7cm^2^ in older ones (6mm diameter). Given the width-length relationship, we considered the diameter of the stem as a proxy to infer the distance from the base of the plant.

To understand the area that each tissue occupies in the cross section of the stems, we measured the radius of the ring formed by the cortex, the phloem, the xylem, and the pith, and calculated the area: *A*= π (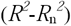), being *A* area, *R*_n_ the radius of each ring tissue, and R the total radius of the stem. We then normalized these areas to the percentage of total cross sectional area occupied, which revealed a decreasing percentage of the cross sectional area occupied by the cortical tissue as the stems increase in diameter (Fig. 2a-c), and an opposite pattern for the xylem tissue. Strikingly, the proportion of the cross-sectional area occupied by the phloem tissue was maintained in all stem diameters evaluated (Fig. 2d). Because the phloem tissue is composed of clusters of sieve tubes and other cells separated by ray parenchyma, we quantified the proportion of sieve tubes per cluster in different stem diameters to infer the total number of tubes. While the proportion of conductive cells (number of sieve tubes/total number of phloem cells in each cluster) was maintained at approximately 30% in all of the stem diameters evaluated, the estimated total number of sieve tubes increased linearly with stem diameter (*r*^2^=0.99; p<0.05; Fig. 2d).

**Fig. 2.**
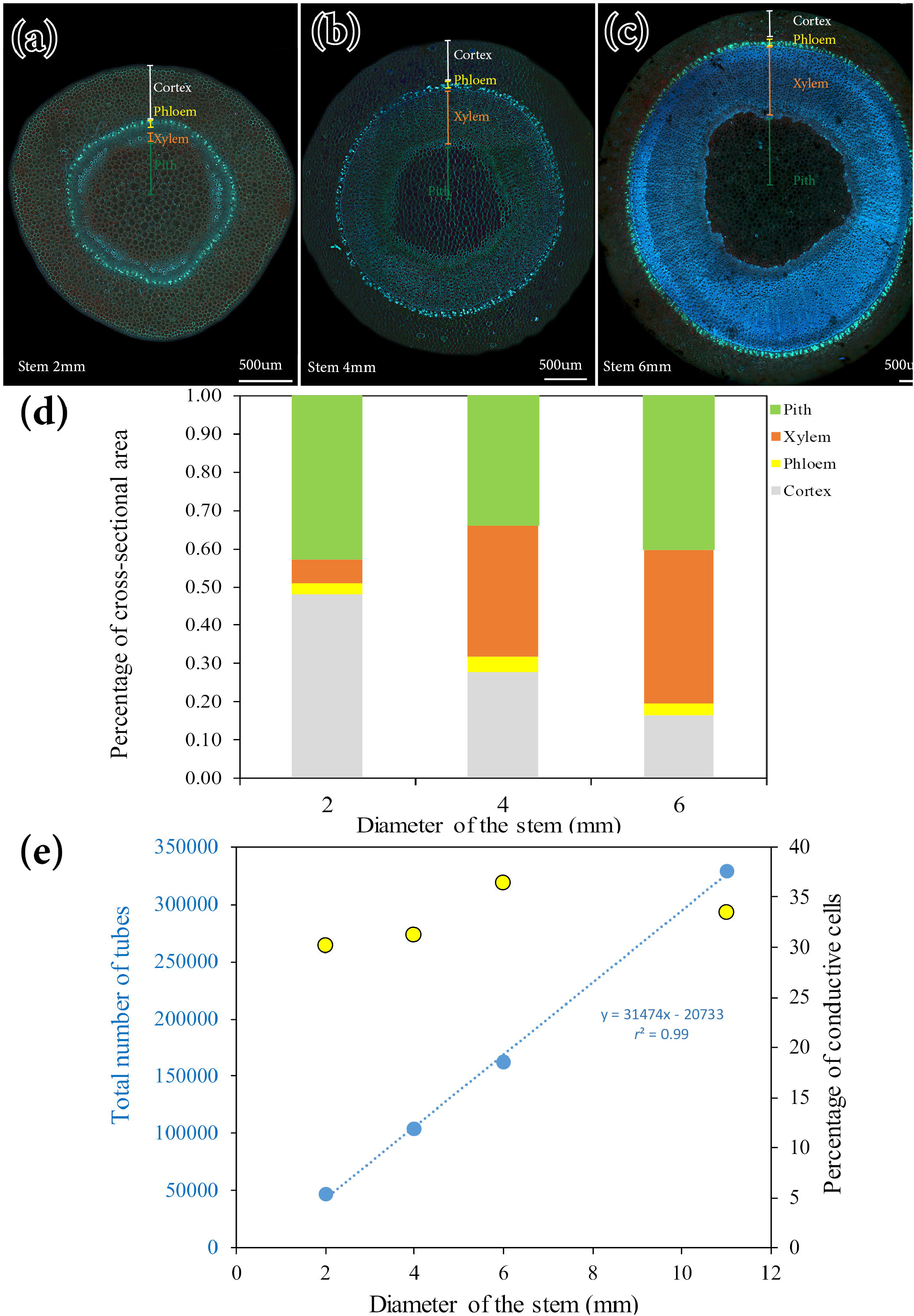
Anatomy of the stems in *Illicium parviflorum*. (a) Stem with primary growth, 2mm diameter. (b) Stem with secondary growth, 4mm diameter. (c) Barky stem, 6mm diameter. (d) Percentage of cross-sectional areas occupied by the cortex (grey), phloem (yellow), xylem (orange), and pith (green) tissues at different developmental stages of the stems based on their diameter. (e) Sieve tube element number per stem cross-sectional area: blue color represents the relationship between the total number of sieve tube elements and stem diameter; yellow dots represent the percentage of sieve tubes (conductive area) with respect to the total number of cells per phloem cluster (the areas of the phloem separated by ray parenchyma). (a-c), 50μm cross sections stained with aniline blue for callose. Bars: 500μm.

The morphology of the individual sieve tube elements further exhibit variation across stem diameters, increasing the number of sieve areas per plate connections between tubes as stem diameters increase (Fig. 3a-c). The length (*l*) and radius (*r*) of individual sieve tube element showed a logarithmic positive relationship with stem diameter (p<0.05; Fig. 3d). Similarly, both the areas of sieve plates as well as their number follow a positive logarithmic relationship with stem diameter (p<0.05; Fig. 3e). The average radius of sieve plate pores in the two large stem diameter classes was 0.22μm^2^ ± 0.005 SE. We were unable to quantify the pore size of smaller diameter stems because the sieve pores were occluded with an amorphous material.

**Fig. 3.**
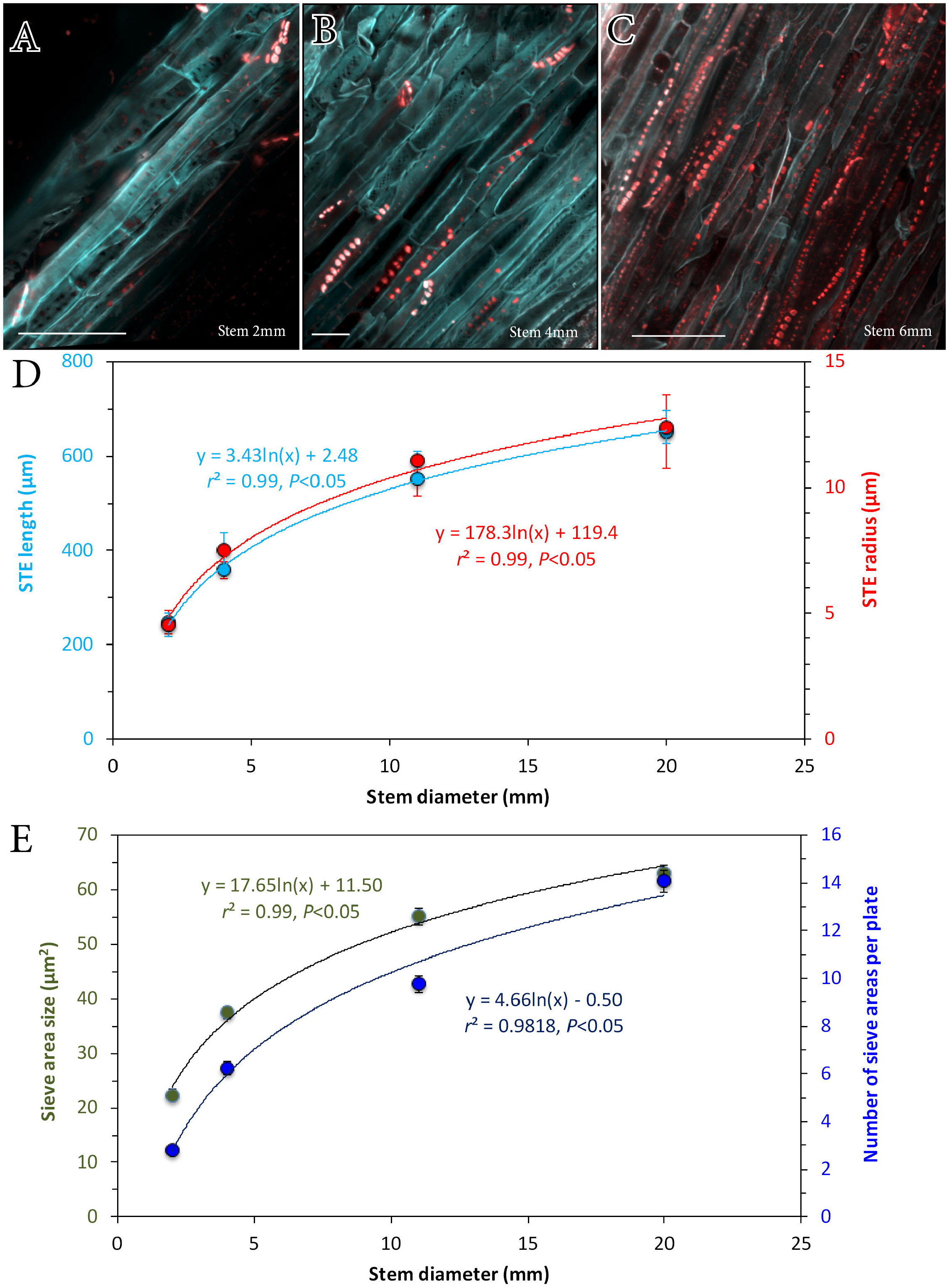
The phloem in the stems of *Illicium parviflorum*. (a) Sieve tube element in a stem with primary growth (2mm diameter) showing cell walls in cyan and sieve plates in red. (b) Sieve tube elements in the 4mm diameter stem. (c) Sieve tube element network in the 6mm diameter stem. (d) Geometrical relationships between sieve tube element length (light blue), radius (red) and diameter of the stem. (e) Relationship between number of sieve areas per end plate (dark blue) and their size (green) across stem diameters. (a-c), longitudinal sections of the stems stained with aniline blue for callose and calcofluor white for cellulose. Logarithmic regression curves at a *p*<0.05, and bars display standard error (SE). Bars: (a), 100μm; (b), 200 μm; (c), 500μm.

### Leaf vascular anatomy in *Illicium parviflorum*

The mature leaves of *I. parviflorum* have reticulate venation; veins range from a small petiole (1cm long on average), midrib through fourth order veins, with higher vein orders difficult to disentangle (Fig. 4a). Cross sections of the major veins in leaves from adult plants revealed that both the phloem and the xylem organize in clusters of conduits, which are axially separated by rays of parenchymatous cells (Fig. 4b), similar to the organization observed in the stem vasculature. While the vascular tissue is mainly composed of xylem and phloem in the petiole, a sheath of thick-walled fibers surrounds the vasculature of the primary and higher order veins in the leaf lamina (Fig. 4c, d). Interestingly, despite the reduction in the total area occupied by the vascular tissues from the petiole towards the tip of the midrib (Fig. S1), the number of phloem and xylem cells in each cluster, as well as their respective lumen areas, is maintained along the entire midrib (Fig. S2). These results suggest that the increase in the conductive areas of the major vein towards the petiole is due solely to the increasing number of vascular clusters, and thus conduits. But it also points to the intercalary growth of the leaf ray parenchyma as a major cause of vein tapering. Although the 1:1 xylem-phloem cell number maintains at the secondary veins, their number and lumen areas are reduced by almost an order of magnitude compared with the major vein. Leaf vasculature is markedly different in seedlings, where the xylem and phloem cells are still differentiating. Between them, several rows of square and thin walled cells likely correspond with cambial tissue (Fig. 4f-h).

**Fig. 4.**
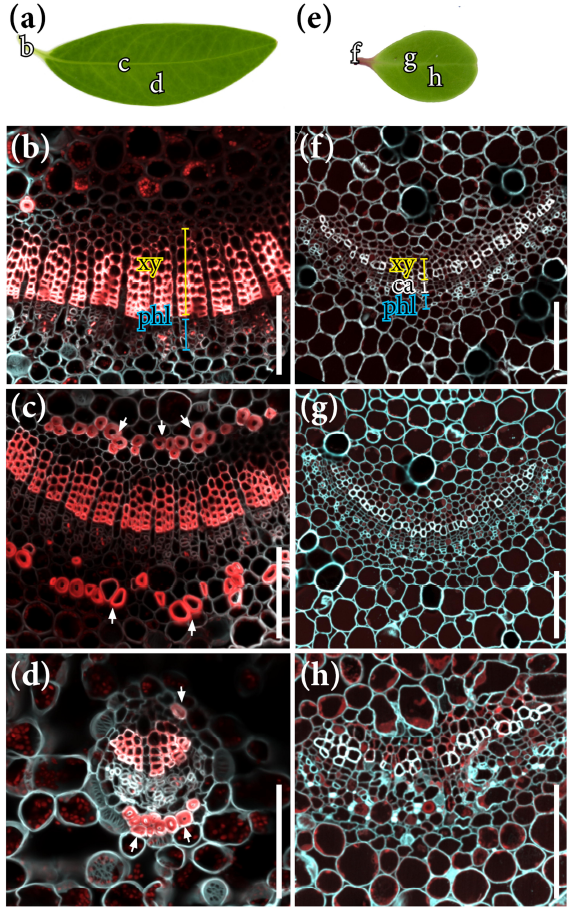
Anatomy of the leaves of *Illicium parviflorum*. (a) Mature leaf showing the adaxial surface and the reticulate venation. (b) Vasculature of the petiole, showing the arrangement of xylem (Xy), and phloem (Phl) in clusters separated by rows of ray parenchyma. (c) Cross section of the midrib showing a similar organization but surrounded by a sheath of sclereid tissue (white arrows). (d) Secondary vein showing reduced xylem and phloem tissue compared with the primary vein. (e) Leaf of a seedling. (f) Petiole showing a differentiating xylem (yellow), several rows of cambial cells (white), and differentiating phloem (blue) tissues. (g) Similar organization in the midrib. (h) Secondary vein. (b-d), (f-h): Cross sections of the stained with aniline and calcofluor white. Xy, xylem; phl, phloem; ca; cambium. Bars: 500μm.

Quantifications of the length and radius of sieve tube elements in leaves of *Illicium parviflorum* reveals that both parameters decrease from the major vein to the higher order veins (Fig. 5a), yet their dimensions are conserved between the midrib and the petiole (Fig. 5b). Unfortunately, detecting the sieve tube elements at fourth vein orders was impossible due to the massive presence of sieve areas along their lateral cell walls (Fig. 5c). Strikingly, the sieve tube elements of the leaves from seedlings were longer and wider compared with mature leaves (Fig. 5d,e).

**Fig. 5.**
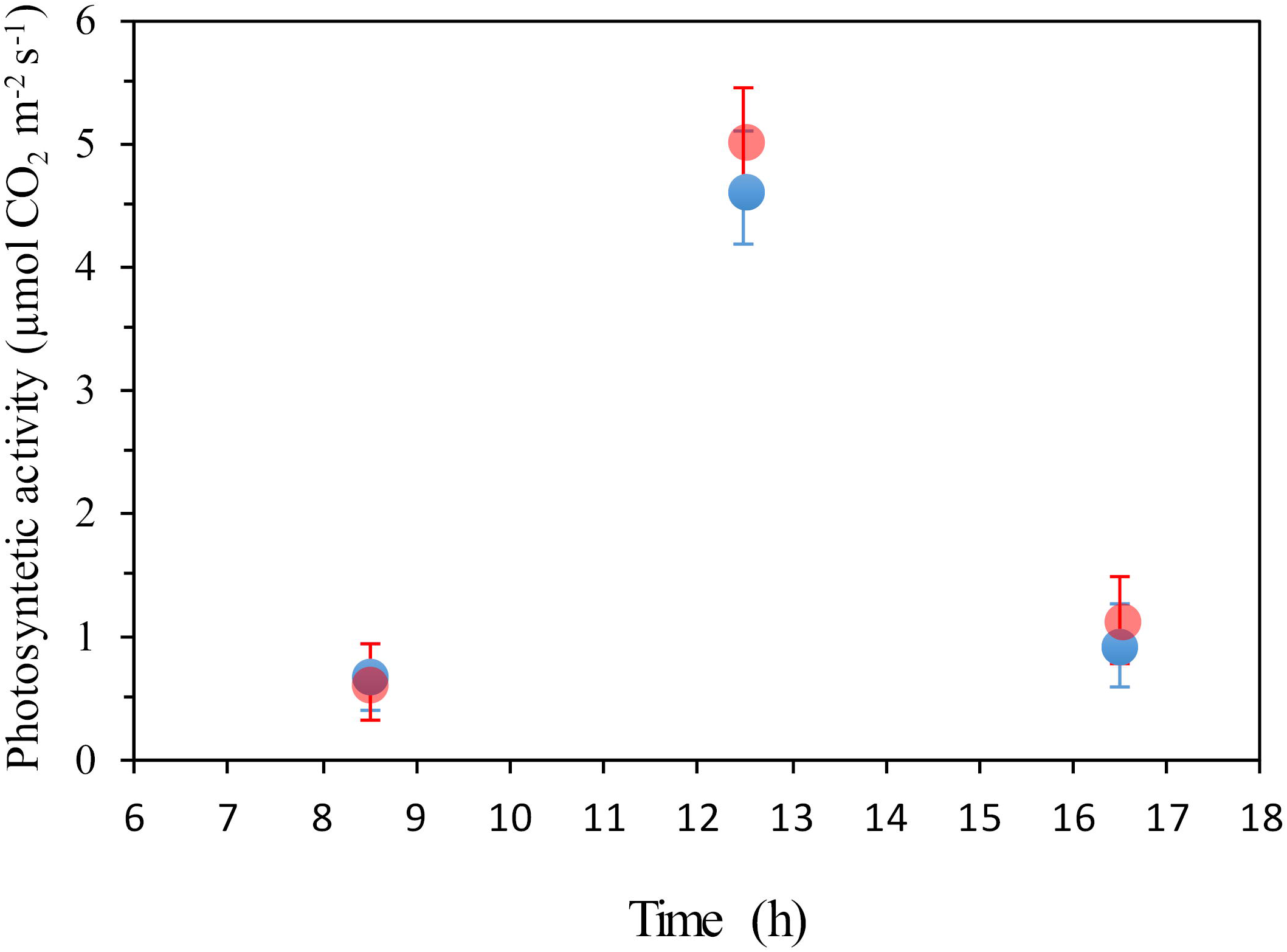
The phloem in the leaves of *Illicium parviflorum*. (a) Scaling relationship between sieve tube element length (blue) and radius (red), across major veins in mature leaves of I. parviflorum. (b) Longitudinal section of the sieve tube elements of the secondary vein of a mature leaf showing the sieve plate connections (arrowheads). (c) Sieve tube element of a fourth order vein in a mature leaf showing sieve plates all along the lateral walls. (d) Relationship between sieve tube element length (blue) and radius (red) in the major veins of seedlings. (e) Longitudinal section of the secondary vein of a leaf from seedlings, with the sieve plate connection (arrowheads) between contiguous tubes. (a,d) bars display standard error (SE); (b,c,e) bars: 500μm.

### Assimilation rates and phloem sap velocity

Photosynthetic activity was not significantly different between adult plants and seedlings of *Illicium parviflorum*. Maximum rates of 4-5 µmol CO_2_ m^−2^ s^−1^ were observed at midday (Fig. 6). To evaluate the velocity of transport through the phloem, we monitored the fluorescence front of a dye tracer along the secondary veins of both adult plants and seedlings (Fig. 7). While this velocity was very low for adult plants, 4.94 ± 1.10 μm s^−1^ (n=3) (compared with other angiosperm species), the veins of seedlings exhibited a velocity that was nearly three times higher (14.44 ± 1.70 μm s^−1^), pointing to a regulation of sap flow velocity through the phloem in *I. parviflorum* plants depending on their developmental stage.

**Fig. 6.**
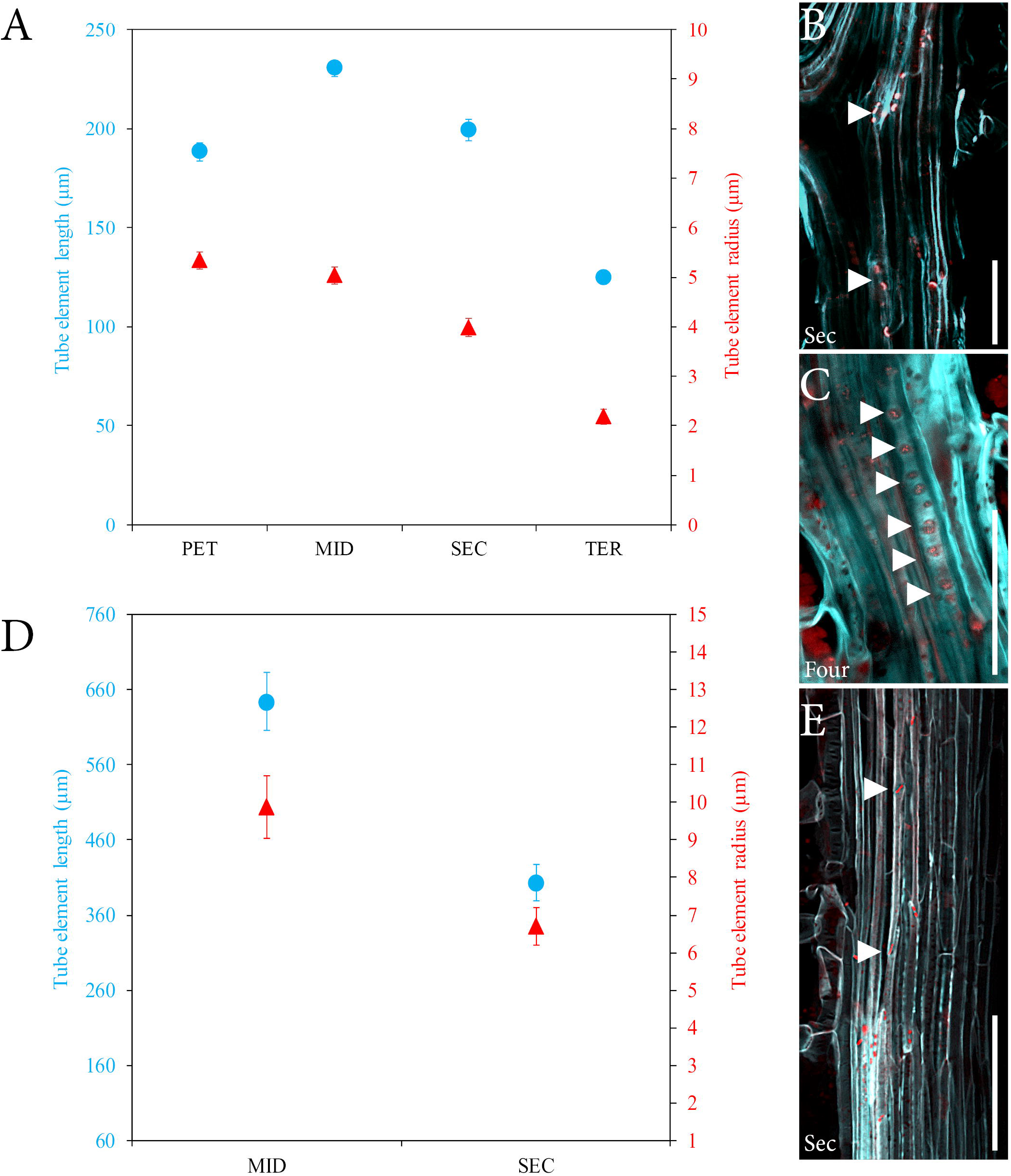
Photosynthetic activity in *Illicium parviflorum*. Comparison between the assimilation rates of adult plants (blue) and seedlings (red) under the same greenhouse conditions at three time points during the day. Error bars (SE) emphasize overlapping averages between both plant types.

**Fig. 7.**
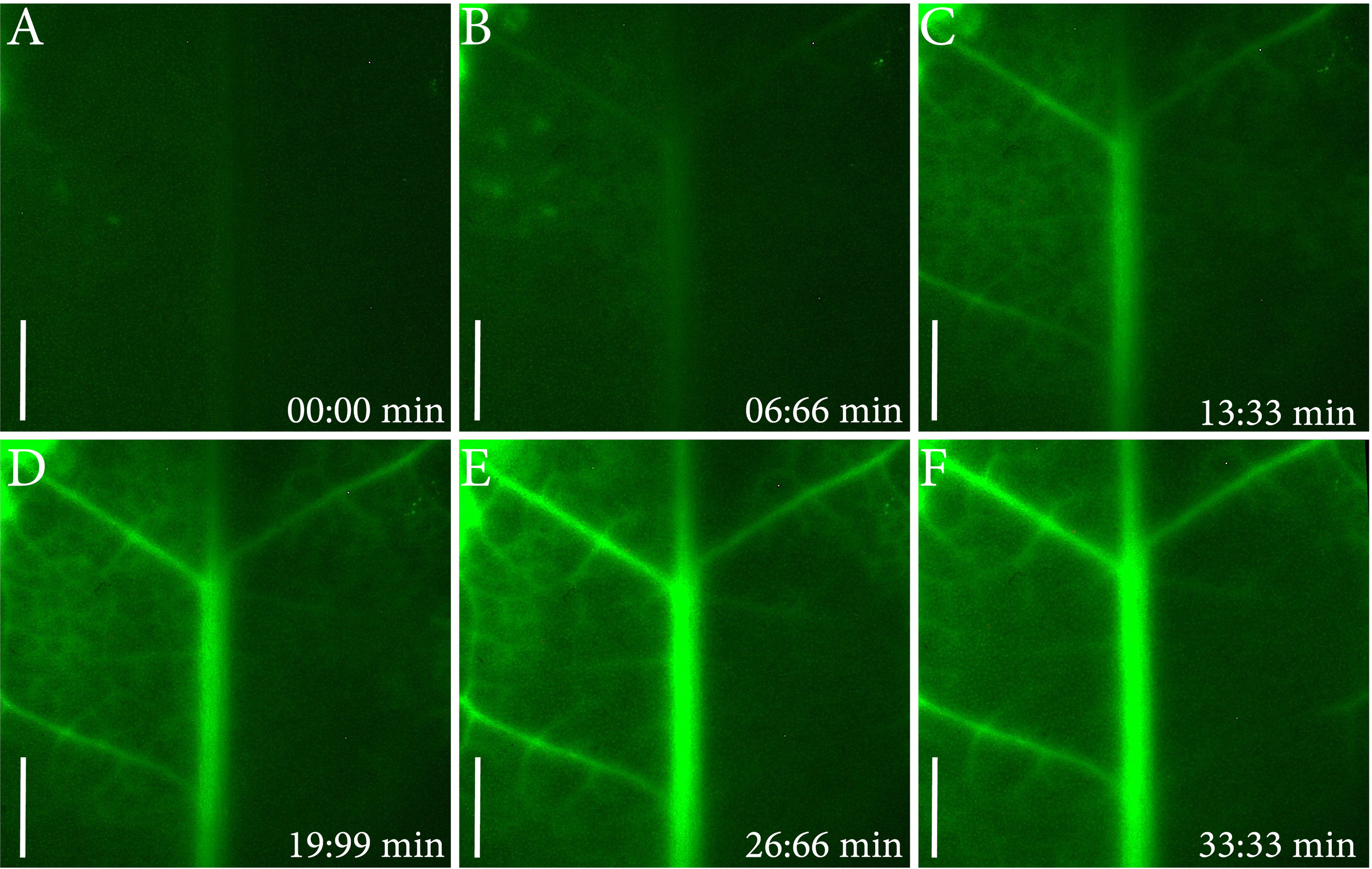
Velocity of the sap within the phloem of *Illicium parviflorum*. Time lapse images of a leaf from a seedling showing advancement of the carboxyfluorescein dye through the phloem. Scale bars: 1000μm.

### Sieve tube hydraulic resistance in stems and leaves

From the collected anatomical data, we computed the hydraulic resistance of sieve tubes from thinner and shorter branches, which have the smallest tubes, to the stems at the base of the shrub, which exhibit the widest diameters and bigger sieve tubes. Sieve tube resistance can be expressed as R_tube_=R_lumen_+R_plate_. R_lumen_ depends on both the geometry of the tubes themselves and the viscosity of the sap flowing within, which we assume here as □=1.7mPa, an estimate obtained in previous measurements (Knoblauch *et al*., 2016). R_plate_ is determined by the number and size of sieve pores (Jensen *et al*., 2012), with pore radius the most influential parameter affecting resistance (see also Jensen *et al*., 2014; Savage *et al*., 2017). Note that because we were unable to measure the pore diameters of the smaller stems, for our calculations we assume that pore size does not vary.

Sieve tube hydraulic resistance per length in *I. parviflorum* has an inverse relationship with the diameter of the stems (Fig. 8a), but the magnitude of this decrease is modest compared to patterns recently reported for trees (Savage *et al*., 2017). To understand the influence of this decreasing resistance on the pressure required to drive sap flow in the leaves, we used the resistor analogy recently applied to trees (Savage *et al*., 2017). This model assumes that the differential pressure across transport length L is proportional to the hydraulic resistance per length of sieve tubes [R (Pa s m^−4^)] and the sap flow rate [Q (m^3^s^−1^)] expressed as Δp/L = QR. We calculated Δp across a transport length (L) for different diameter stems, assuming a constant linear flow velocity (*U*), and the average sieve element radius (*r*) and length (*l*), and our estimate of sieve tube resistance [R_tube_ (Pa s m^−3^)]:

**Fig. 8.**
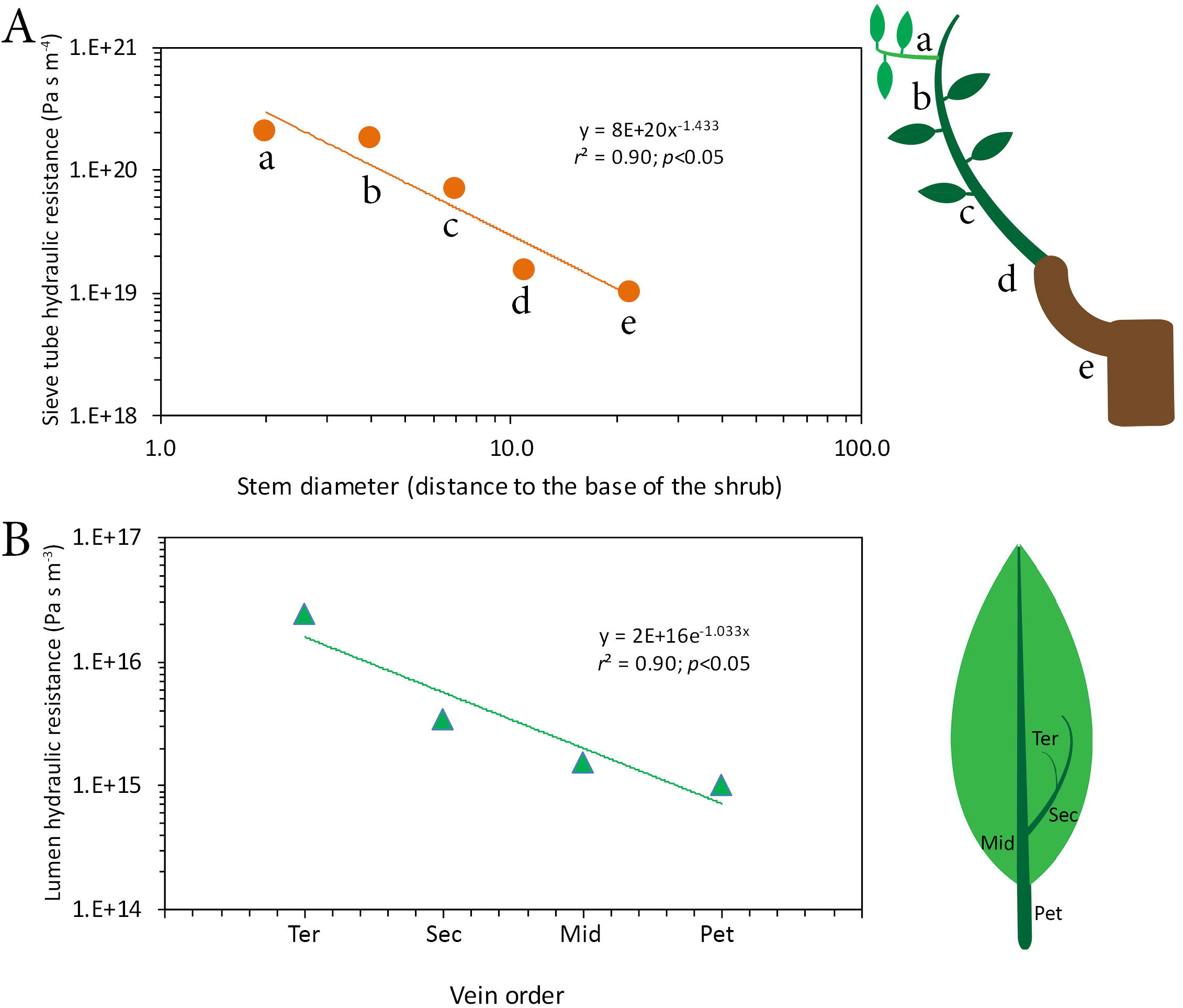
Sieve tube hydraulic resistance in the aerial parts of *Illicium parviflorum*. (a) Inverse relationship between the sieve element resistances per length at each stem diameter (related with the distance to the base of the stem). (b) Sieve tube lumen resistance in the major veins of the mature leaves of *Illicium parviflorum*.

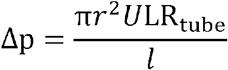

Assuming the phloem sap velocity measured in mature plants (5 µm s^−1^), the differential pressure required to transport sap in a 1-m-long branch is very low (0.05MPa for the smallest branch diameters). Thus, with a fixed viscosity and relatively low pressures, the small branches of *Illicium parviflorum* could be much longer without compromising carbohydrate transport. The higher phloem sap velocity measured in seedlings implies that the pressure required to transport the same distance would increase, yet the estimated pressure difference (0.16 MPa) is within the range of most measured angiosperms. It has to be noted that seedlings, similar to younger branches, rarely reach lengths longer than 0.05m without widening their stems.

Using the geometrical parameters of the sieve tube elements from leaves, we evaluated the lumen hydraulic resistance in different vein orders (Fig. 8b). As in the stems, phloem hydraulic resistance decreased by over an order of magnitude from tertiary veins to the petiole.

## DISCUSSION

### Scaling of phloem geometry in the stems of *Illicium parviflorum*

The current work shows that the increase in stem diameter in *I. parviflorum* towards the base of the shrub correlates with a developmental scaling of sieve tube structure. Comparable values of length and width of the sieve tubes have been only recently reported in angiosperm trees, pointing to a conserved mechanism associated with attainment of an arborescent growth form (Liesche *et al*., 2017; Savage *et al*., 2017). However, pore sizes of the sieve plates were substantially smaller in *Illicium*. The pore areas of the sieve plates have been estimated as the most influential parameter affecting sap flow resistance within sieve tube (Esau *et al*., 1962; Mullendore *et al*., 2010; Jensen *et al*., 2012). Thus, the decreased resistance of the phloem towards the base of the shrubs in *Illicium* is mainly caused by the scaling of geometrical properties of the sieve tube elements, but especially the increasing number of sieve areas per plate in the sieve tube end walls. If this holds true for other basal angiosperms, small pore size could be an ancestral feature of woody flowering plants related with a high phloem resistance, which is consistent the high xylem resistance associated with the low density of vessels previously reported in the wood (Feild *et al*., 2003a; 2004; Feild *et al*., 2009).

Interestingly, the proportional cross-sectional area of the phloem in different stem diameters of *I. parviflorum* is conserved, contrasting with the reduction of cortical tissue, as well as the increase in xylem tissue. The scaling of xylem elements towards the base of woody angiosperms, as well as their increase in number have been widely studied (Sevanto *et al*., 2011; Hölttä *et al*., 2006; 2009; 2013; Petit & Crivellaro, 2014; Diaz-Espejo & Hernandez-Santana, 2017). This is because anatomical quantification of vessels in cross section is straightforward, whereas counting the sieve tube elements is more challenging. Due to the paired structure-function relationship between the phloem and the xylem (reviewed in Savage *et al*., 2016; Seleznyova & Hanan, 2018), understanding hydraulic transport in woody organisms requires a better understanding of sieve tube numbers along the plant, so far missing in woody lineages. Previous sieve tube number estimations were based on single point measurements in the stems, such as the 2/3 sieve tube proportion (number of sieve tubes relative to total phloem cells) inferred by Münch (Münch, 1930), which was later challenged by quantifications of sieve tubes in the stems of other angiosperm trees, such as the 12-26% in six *Cassia* species (Ghouse & Jamal, 1979), 17-35% in members of the Myrtaceae family (Ghouse *et al*., 1976), 54-74% in *Sterculia tragantha* and *Bombax bounopozense* (Lawton & Canny, 1970), or 11-59% in leguminous trees (Khan *et al*., 1992). These single point measurements offer only a partial picture (see also Canny, 1973). Here, we use serial sections to estimate the total number of sieve tubes per cross sectional area of the stem at different positions along the plant. While the proportion of conductive elements per phloem cluster (30%), are maintained in all stem diameters in *Illicium parviflorum*, consistent with the similar proportion of vessels in the xylem (Feild *et al*., 2003a), the total number of conductive tubes increases linearly with stem diameter. Interestingly, in stems with primary growth, the average leaf area supplying photosynthates doubles from thinner to thicker stems and the estimated number of sieve tubes follows a similar trend. The lack of sieve tube number quantification across different stem diameters in the majority of woody angiosperms hampers a comparative framework with our data, and even though future works will elucidate whether this pattern shows consistency across woody plants, our results point to a critical spatial-temporal regulation of sieve tube number and size in stems.

### Phloem scaling in leaf veins of *I. parviflorum*

Unlike in the stems, mature leaves of *Illicium parviflorum* show a similar number of vascular elements of both xylem and the phloem, a relationship that appears to be conserved across vein orders. These results are consistent with the idea of a physiological dependence between xylem and phloem (Zwieniecki *et al*., 2004; Sevanto *et al*., 2011; Hölttä *et al*., 2013), but contrast with recent measurements in herbaceous plants (Ray & Jones, 2018). Additionally, the proportion of sieve tubes examined in the petioles of herbaceous species vary substantially, from the roughly 30% of sieve tubes in *Beta vulgaris* (Geiger *et al*., 1969), to 17% and 23% of *Cucurbita* species and potato tubers respectively (Crafts, 1931; 1933). While this has consequences for mass transfer from the leaves to the stems, the 1:1 relationship between the phloem and the xylem in the leaves of *I. parviflorum* implies that the increasing total conductive area of the xylem and phloem in the major veins results from a higher number of conduits towards the petiole, similar to the described reduction in phloem conduit number in the singled veined needles of pines (Ronellenfitsch *et al*., 2015). In addition, different vein hierarchies in the leaves of *I. parviflorum* display a scaling relationship of sieve tubes that is similar to what was recently reported for the reticulate veined leaves of poplar (Carvalho *et al*., 2017a) and the dichotomously veined *Ginkgo* (Carvalho *et al*., 2017b). However, the sieve tube elements are shorter and thinner at the petiole in *Illicium parviflorum* leaves than within the leaf veins. Shorter sieve tubes in the phloem of the petiole may have implications for the regulation of pressure within the sieve tubes in leaves, especially at the times of maximum turgidity (i.e. maximum sugar export rates). We observed a higher number of sieve areas in the plates connecting the sieve tubes in the petiole, thus this feature may attenuate at least in part a putative pressure increase (Carvalho *et al*., 2018). It would be desirable to obtain pore size in the sieve tubes of different vein orders and therefore check whether it concords with the hydraulic models for energy conservation, such as the da Vinci’s or Murray’s rules (Murray, 1926; Richter, 1980; McCulloh *et al*., 2003). Nevertheless, the role of the petiole regulating sugar export from leaves has not been extensively explored across angiosperms and requires further attention (Grimm *et al*., 1997; Ray & Jones, 2018).

The directional flow in the sieve tubes of the major veins in *Illicium* leaves contrasts with the anatomy of the sieve tubes in the minor veins, where numerous sieve areas pervade their lateral walls. This is in line with the idea of a division of function between the minor veins, which mainly work as sugar loaders, and major veins, where directional transport occurs within leaves (Russin & Evert, 1985; Turgeon, 2006; Carvalho *et al*., 2017a, 2018). A high number of symplasmic connections typically associate with passive sugar loading in the minor veins (van Bel *et al*., 1992; Turgeon, 1996; Gamalei *et al*, 2000; Rennie & Turgeon, 2009; Turgeon, 2010; Davidson *et al*., 2011; Zhang *et al*., 2014), but the heterogeneity of the species evaluated leave this question still unresolved (Slewinski *et al*., 2013). So far, sugar (radio) labeling appears as the most reliable measure of loading type, which has yet to be applied to *Illicium parviflorum* leaves. As a member of the basal grade of flowering plants, *Illicium* could likely fit with the previously hypothesized passive sugar loading in woody angiosperms of the understory (Gamalei, 1989; 1991). However, this feature appears to be labile among the extant members of the basal angiosperm grade, such as the active loading reported for *Amborella* (Turgeon & Medville, 2011; Comtet *et al*., 2017).

### Developmental flexibility and carbon limitation at the sources in the understory

Our estimations of the turgor pressures required to drive sap transport in *I. parviflorum* are within the range of osmotic values for phloem sap reported for a wide range of species (Jensen *et al*., 2013). These measurements thus support the validity of the Münch hypothesis (Münch, 1930) as the mechanism of sugar transport in understory shrubs, consistent with models of phloem transport in both angiosperm and gymnosperm trees (Thompson & Holbrook, 2003; Liesche *et al*., 2015; Jyske & Holtta, 2015; De Schepper *et al*., 2013; Comtet *et al*., 2017). However, our calculations indicate that even without increases in stem diameter, *I. parviflorum* branches could reach significant lengths without compromising phloem transport.

In addition, our evaluations of carbon low assimilation rates in greenhouse conditions are consistent with the low *in situ* measurements previously reported in the field (Feild *et al*., 2003a,b; 2004; Feild *et al*., 2009). Interestingly, the size of sieve tube elements, both diameter and length, in the major veins of the leaves of seedlings are approximately three times larger than in mature leaves. In parallel, the measured velocity of sap in the seedlings is threefold higher than that of adult leaves, suggesting a functional flexibility of the phloem through ontogeny. Faster transport rates through the phloem of saplings would imply a more efficient transporting of sugars during rapid sun flecks, which account for one third of the total photosynthesis in the tropical understory (Pearcy, 1987). While phloem velocity of seedlings and adult plants has only been evaluated in a single herbaceous species (Savage *et al*., 2013), they observed variable phloem sap velocity in seedlings depending on the bundle and developmental stage. However, whether this developmental dynamism is consistent across flowering plants requires further investigations.

### Phloem dynamics and the ecophysiology of early angiosperms

Despite the understory origins of woody angiosperms, organismal scaling of sieve tube elements of the phloem appears as a conserved mechanism that predates the origin of woody flowering plants, and likely seed plants in general (see Woodruff *et al*., 2004; Liesche *et al*., 2015; Liesche, 2017). Angiosperm colonization of the vertical niche implied a number of structural innovations that led to a higher functional efficiency and thus productivity (Koch *et al*., 2004; Savage *et al*., 2017; Gleason *et al*., 2018). However, our knowledge on phloem functioning in the extant lineages of the basal angiosperm grade is still in its infancy. We hereby reported for the first time a detailed evaluation of phloem anatomy and physiology in a member of the basal-grade of angiosperms, offering a developmental angle that may help to explain the plasticity of the phloem in the light-limited environments where these lineages likely evolved. With respect to the phloem of adult plants, the major structural difference between the sieve tubes of *I. parviflorum* and those found in angiosperm trees is their much smaller sieve plate pores. This suggests that evolutionary changes in sieve tube structure could have critically influenced the structure and functioning of forest ecosystems.

Extant angiosperm taxa composing the basal-branch lineages of flowering plants are no more than 220 species, yet the spectrum of their life forms range from aquatic herbaceous (the whole Nymphaeales clade), to shrubs and lianas of the tropical understory in both the monotypic Amborellales and the Austrobaileyales. From a physiological perspective, inferring the evolution of physiological performance during angiosperm radiation has typically been carried out from a single organ perspective, such as leaves (Brodribb & Feild, 2010) or vascular traits of the xylem in mature plants (Feild & Arens, 2005, 2007). However, broader reconstruction of ancestral ecophysiological traits further requires organismal and developmental approaches, especially from the perspective of the understudied phloem tissue. This is particularly relevant in plants that acquired arborescent forms, since their size requires efficient internal transport tissues and they play an outsize role in structuring forest ecosystems. We propose that developmental and organismal heterogeneity of sieve tube elements across extant member of the basal angiosperm grade (see Behnke, 1986) are key elements for the reconstruction of traits that compose different strata in ancestral and contemporary forests. Thus, suites of functional vascular traits including perforation plates of the xylem ((Feild and Wilson, 2012), as well as sieve plates of the phloem, may have been selected during early angiosperm evolution, allowing woody flowering plants to extend beyond their understory origins.

## Acknowledgements

This work was funded by the National Science Foundation IOS 1456682 research grant to NMH. We are very grateful to William E. (Ned) Friedman for kindly sharing plant material and lab facilities, as well as the staff of the Arnold Arboretum of Harvard University for continuous support. We are thankful to Yongjiang Zhang for helping with LI-COR measurements, as well as Jessica Gersony and Kasia Zieminska for their thoughtful feedback on the manuscript.

## Author contributions

NMH conceived the project, supervised the experiments and data, and wrote the manuscript. JML designed and performed the experiments, analyzed the data, and wrote the manuscript.

## Supporting information Figures S1 and S2

Vascular tissues in the major veins of mature leaves and seedlings in *Illicium parviflorum.*

**Supporting information Figure S1.** Cross-sectional area of the vascular tissues along three areas of the major veins of mature leaves in *Illicium parviflorum*: the petiole, the midrib and the tip of the mayor vein. Letters over bars (SE) show significant differences between the areas at each point using the Tukey test for analysis of variance at a p<0.05.

**Supporting information Figure S2.** Linear correlations between the total number of phloem cells and cambial cells (black) per vascular cluster, as well as the number of phloem cells and xylem cells (grey) in either mature leaves (top) and seedlings (bottom).

